# Visualizing the dynamics of exported bacterial proteins with the chemogenetic fluorescent reporter FAST

**DOI:** 10.1101/2020.05.12.090142

**Authors:** Yankel Chekli, Caroline Peron-Cane, Dario Dell’Arciprete, Jean-François Allemand, Chenge Li, Jean-Marc Ghigo, Arnaud Gautier, Alice Lebreton, Nicolas Desprat, Christophe Beloin

## Abstract

Bacterial proteins exported to the cell surface play key cellular functions. However, despite the interest to study the localization of surface proteins such as adhesins, transporters or hydrolases, monitoring their dynamics in live imaging remains challenging, due to the limited availability of fluorescent probes remaining functional after secretion. In this work, we used the *Escherichia coli* intimin and the *Listeria monocytogenes* InlB invasin as surface exposed scaffolds fused with the recently developed chemogenetic fluorescent reporter protein FAST. Using both membrane permeant (HBR-3,5DM) and non-permeant (HBRAA-3E) fluorogens that fluoresce upon binding to FAST, we demonstrated that fully functional FAST can be exposed at the cell surface and specifically tagged on the external side of the bacterial envelop in both diderm and monoderm bacteria. Our work opens new avenues to study of the organization and dynamics of the bacterial cell surface proteins.

## Introduction

The study of protein localization dynamics using fluorescent reporters has led to major insights into many biological processes. For instance, use of fluorescent reporters enabled to show that MreB, the actin-like protein found in bacteria, spatially dictates the subcellular sites of cell wall synthesis ^1,2^. Super folder-GFP fusions with the MinC, D and E proteins, allowed the observation of their oscillation from pole to pole clarifying the role of the Min system in the division septum positioning ^3,4^. Nevertheless, whilst most studies on protein dynamics have been performed on either periplasmic or cytoplasmic proteins, visualizing protein dynamics on the outer bacterial envelope has remained a challenge. Indeed, GFP or GFP-like fluorescent reporters are either blocked before export, or denatured after crossing the bacterial secretion machineries.

To tackle these issues, alternatives to GFP-derived methods have been developed ^5^. In *Escherichia coli*, for instance, fluorescent inactivated colicins have been shown to interact specifically with outer-membrane receptors but their use is restricted to the colicin cognate receptors ^6^. Site-specific covalent labeling of cysteines using thiol-reactive maleimide dyes ^7,8^ or Fluorescein-based biarsenical dyes (FlAsH) ^9^ have been used to label cysteines from polymeric protein structures such as flagellar filaments and type IV pili or bacterial effectors. However, cysteine labeling can be toxic and suffers from a low signal-to-noise ratio, making it more suitable for the study of proteins with repeated motifs. The fluorescence of the Light Oxygen or Voltage (LOV) domain from *Arabidopsis thaliana* phototropin 2 has also been used to tag a flagellin protein in *C. difficile* ^10^ or IpaB in *Shigella* ^11^ but the near-UV excitation of the fluorophore raises issues of phototoxicity during continuous imaging. More recently, the SpyTag/SpyCatcher system based on the covalent tethering of a recombinant reporting domain ^12^, has been used to label a truncated intimin exposed on the outer membrane of *E. coli* ^13^. However, this method is limited by the duration of the labeling process itself, and the protocol involves washing steps to remove the unwanted background caused by the fluorescent dye in the medium, which prevents real-time observation of *de-novo* synthesized proteins.

Recently, a 14 kDa monomeric protein derived from the photoactive yellow protein from *Halorhodospira halophila*, called FAST, was developed as a new chemogenetic fluorescent reporting system ^14^. FAST has been engineered to reversibly bind a fluorogen called hydroxybenzylidene rhodanine (HBR). HBR and its analogues are non-fluorescent by themselves, but their interaction with FAST activates their fluorescence. This property prevents any nonspecific fluorescence even when the fluorogen is present in large excess, allowing to accurately localize FAST-tagged proteins ^15^. FAST has already been successfully used in different organisms such as mammalian cells, zebrafish, yeast and bacteria ^14^. Moreover, this system can be used in anaerobic conditions since fluorescence only depends on the interaction between FAST and the fluorogen. This feature allowed the study of anaerobic organism like *Clostridium* or the study of bacteria in anaerobic environments such as biofilms ^16,17^.

In this study, we showed that FAST can be exposed on the surface of gram-negative (*Escherichia coli*) and gram-positive (*Listeria monocytogenes*) model bacteria. FAST is still able to bind its fluorogenic ligand after its secretion to the surface of bacteria, thereby allowing fluorescence imaging. To characterize the cell uptake of the non-permeant fluorogen HBRAA-3E^18^ through the outer and inner membranes, we localized FAST constructs in the different *E. coli* cell compartments. Finally, we showed that FAST is suitable for monitoring the dynamics of tagged proteins within growing microcolonies for several hours. Overall, these results demonstrate the versatility of the FAST fluorescent reporter system to label proteins exposed on the surface of bacteria and to follow their localization in living cells during colony growth.

## Results and Discussion

### FAST can be exported to the cell surface of *E. coli*

To assess whether FAST can be exported and exposed at the cell surface of *E. coli* we used an anchoring module based on the scaffold of the intimin protein, an outer membrane protein expressed by Enterohemorrhagic *E. coli* (EHEC) and Enteropathogenic *E. coli* (EPEC), and required for intimate attachment to the host cell. The intimin scaffold was shown to efficiently display nanobodies at the *E. coli* cell surface ^19,20^. It is composed of the so-called Neae N-terminal fragment of intimin that encompass a N-terminal signal peptide, a periplasmic LysM domain (allowing binding to peptidoglycan), a β-barrel domain that allows the anchoring of intimin in the outer membrane of bacteria and two Ig-like domains (D00-D0) (Fig. 1a). In the plasmid pNeae2 ^21^ three different tags (E-tag, His-tag and Myc-tag) have been fused in frame with the C-terminal end of the D0 domain. We introduced the DNA encoding the FAST polypeptide after the three tags. In this configuration, D00-D0, the three tags and FAST were expected to be exposed in this order at the cell surface of *E. coli* (Fig. 1a & 1e). The construction is under the control of the IPTG-inducible p*lac* promoter and induction of the expression of the intimin-FAST encoding construct resulted in the production of full-length intimin-D00-D0-tags-FAST (expected size = 86 kDa) as detected using an anti-E-tag antiserum (Fig. 2a). Some extra bands were also observed suggesting the formation of higher order species as well as discrete degradation of the chimeric protein. When using the same antiserum and immunofluorescence on intact cells, in presence of IPTG, we detected a clear fluorescent signal at the periphery of *E. coli* cells (Fig. 2b), thus suggesting that the chimeric construction was properly exported to the outer membrane and exposed at the cell surface by the β-domain of the intimin.

**Figure 1.**
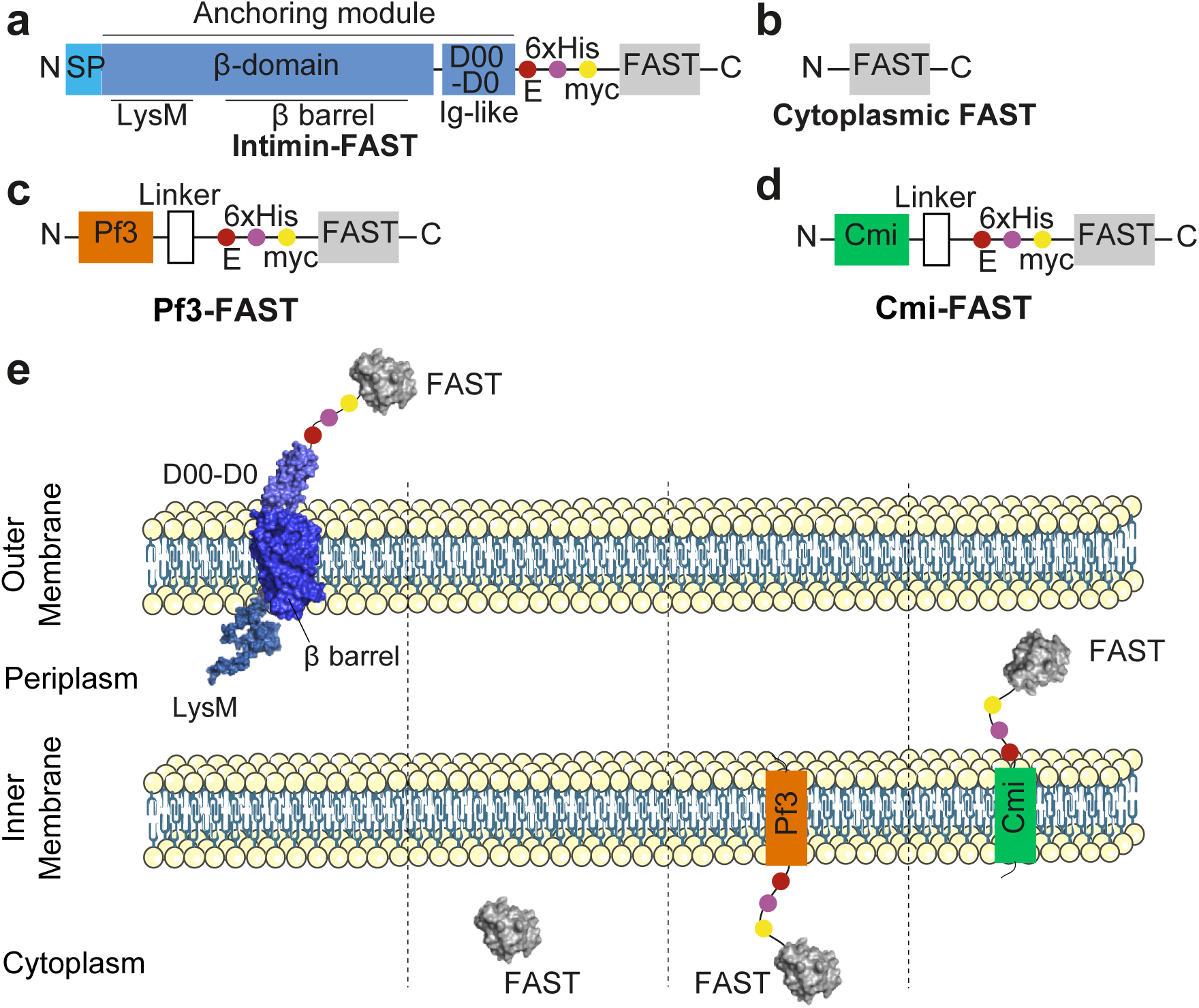
Schematic representation of the different FAST constructs used in this study. FAST was fused to different scaffolds to allow its presentation as an *E. coli* cell-surface protein through the anchoring domain of the intimin (**a**, **e**), a free cytoplasmic protein (**b**, **e**), an inner-membrane attached cytoplasmic protein through the use of Pf3 transmembrane domain (**c**, **e**) or an inner-membrane atttached periplasmic protein through the use of Cmi transmembrane domain (**d**, **e**).

**Figure 2.**
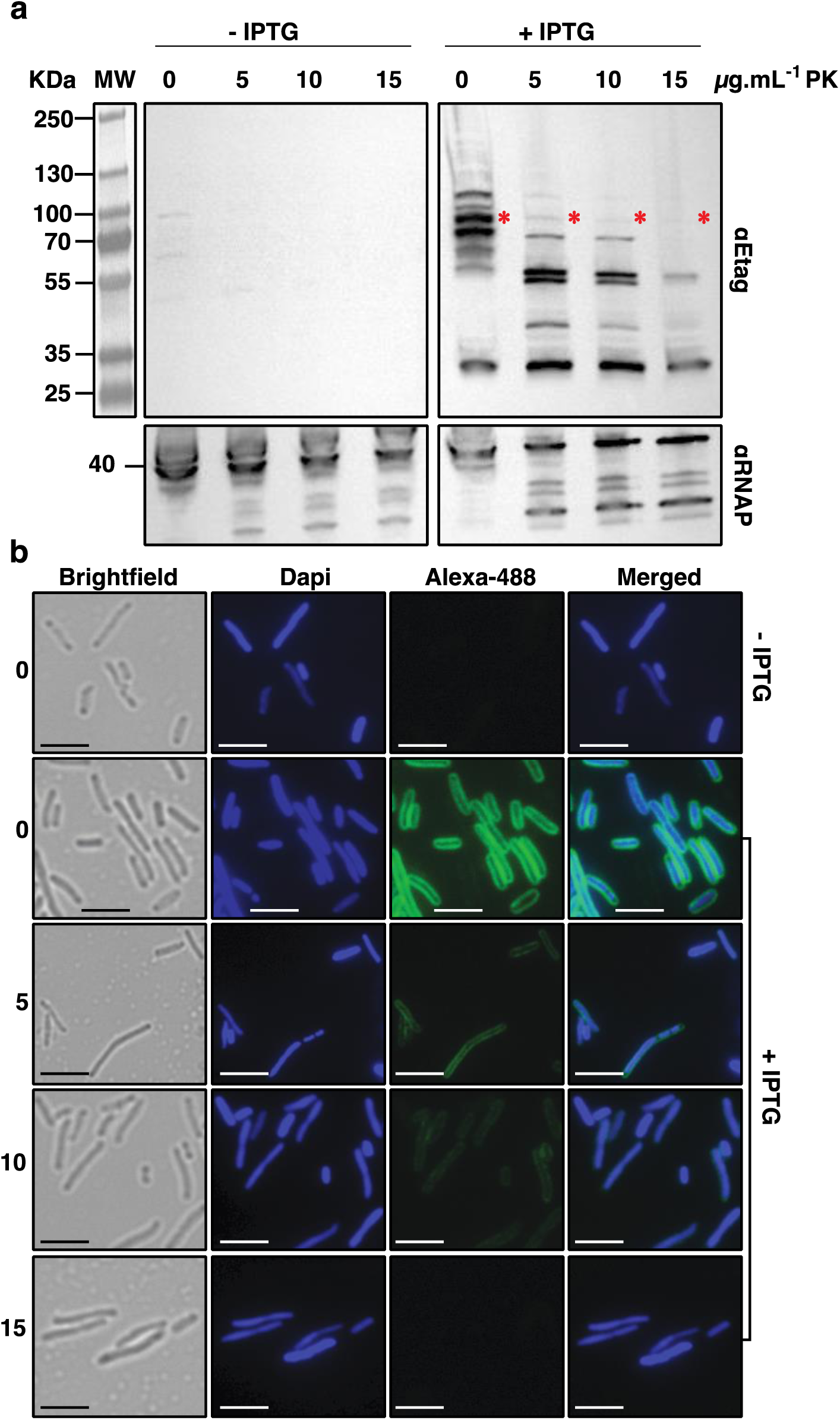
The chimeric Intimin-FAST protein is exposed at *E. coli* cell surface. *E. coli* MG1655 harbouring pNeae2-FAST (expressing intimin-FAST construct) cells were grown overnight in absence or presence of 0.25 mM IPTG to induce the production of the chimeric Intimin-FAST protein. Bacteria were treated with 0, 5, 10 or 15 µg.mL^-1^ of proteinase K for 15 minutes at 40°C before **a:** Western blot of whole-cell protein extracts and revelation with a rabbit anti-E-tag primary antibody and a HRP-linked anti-rabbit secondary antibody, and **b:** Immunofluorescence on whole cells using a rabbit anti-E-tag primary antibody and an anti-rabbit Alexa 488 conjugated secondary antibody. All images are displayed with the same constrast. In **a**, red stars indicate the position of the full length intimin-FAST. In **b**, scale bars = 2 µm.

To confirm that the chimeric protein was indeed transported to the cell surface we assessed its sensitivity to degradation by proteinase K. We performed immunofluorescence and western-blot experiments using cells untreated or treated with increasing amounts of proteinase K (0, 5, 10 or 15 µg.mL^-1^) in order to digest the protein exposed at their surface. The experiment was also performed on a control strain that did not express the chimeric protein. When adding increasing concentration of proteinase K, high molecular weight bands detected in western blot, including the band corresponding to the full-length protein (i.e. the protein exposed at the surface), disappeared. Only low molecular weight bands remained, probably corresponding to protein fragments that are not exposed at the bacterial surface (Fig. 2a). Accordingly, immunofluorescence signals detected at the cell surface for the intimin-FAST chimeric protein progressively disappeared with increasing proteinase K concentration (Fig. 2b). Quantification of the mean fluorescence intensity of cells for each condition confirmed the results shown by western blot and immunofluorescence (Supplementary Fig. S1). Indeed, the use of 5 and 10 µg.mL^-1^ of proteinase K led to a significant 2.5 fold decrease of quantified fluorescence compared to the negative control. This decrease was even more striking when using 15µg.mL^-1^ of proteinase K, with an 18-fold decrease observed between the two conditions. Altogether these results show that FAST was accessible to proteinase K and thus exposed on the surface of the bacteria when fused to the intimin translocator domain.

### FAST can be used to fluorescently tag the beta domain of intimin

While detection of intimin-FAST chimeric constructs on the cell surface by anti-E-tag immunofluorescence demonstrated FAST export, the demonstration of the functionality of FAST requires the use of fluorescence microscopy in presence of FAST fluorogenic ligands in the extracellular medium. Upon binding to FAST, the membrane permeant fluorogen HBR-3,5DM becomes fluorescent (Fig. 3a). We used total internal reflection fluorescence (TIRF) microscopy to visualize the distribution of the intimin ß-domain on the bacterial surface (Fig. 3b). For observations, cells were sandwiched between a glass coverslip and a LB agar pad containing the HBR-3,5DM fluorogen. Brightfield images were obtained from correlation imaging ^22^. Cells expressing the intimin-FAST construct were fluorescent when IPTG and HBR-3,5DM were added to the LB agar pad (Fig. 3c). Due to leakage of the p*lac* promoter, cells containing the intimin-FAST construct were still weakly fluorescent even in the absence of IPTG (Fig. 3c). As expected, only a very weak signal, reflecting the non-specific fluorescent background, was detected for bacteria that did not harbor the intimin-FAST construct in presence of HBR-3,5DM (Fig. 3c). These results thus indicated that functional intimin-FAST fusions could be specifically detected using the HBR-3,5DM fluorogen. Of note, compared to immunofluorescence experiments, the evanescent wave generated by TIRF microscopy excites fluorescence only over the first 200 nm above the coverslip. Consequently, the contour of the membrane cannot be distinguished (Fig. 3b), and fluorescence is only excited in the vicinity of the coverslip (Fig. 3c).

**Figure 3.**
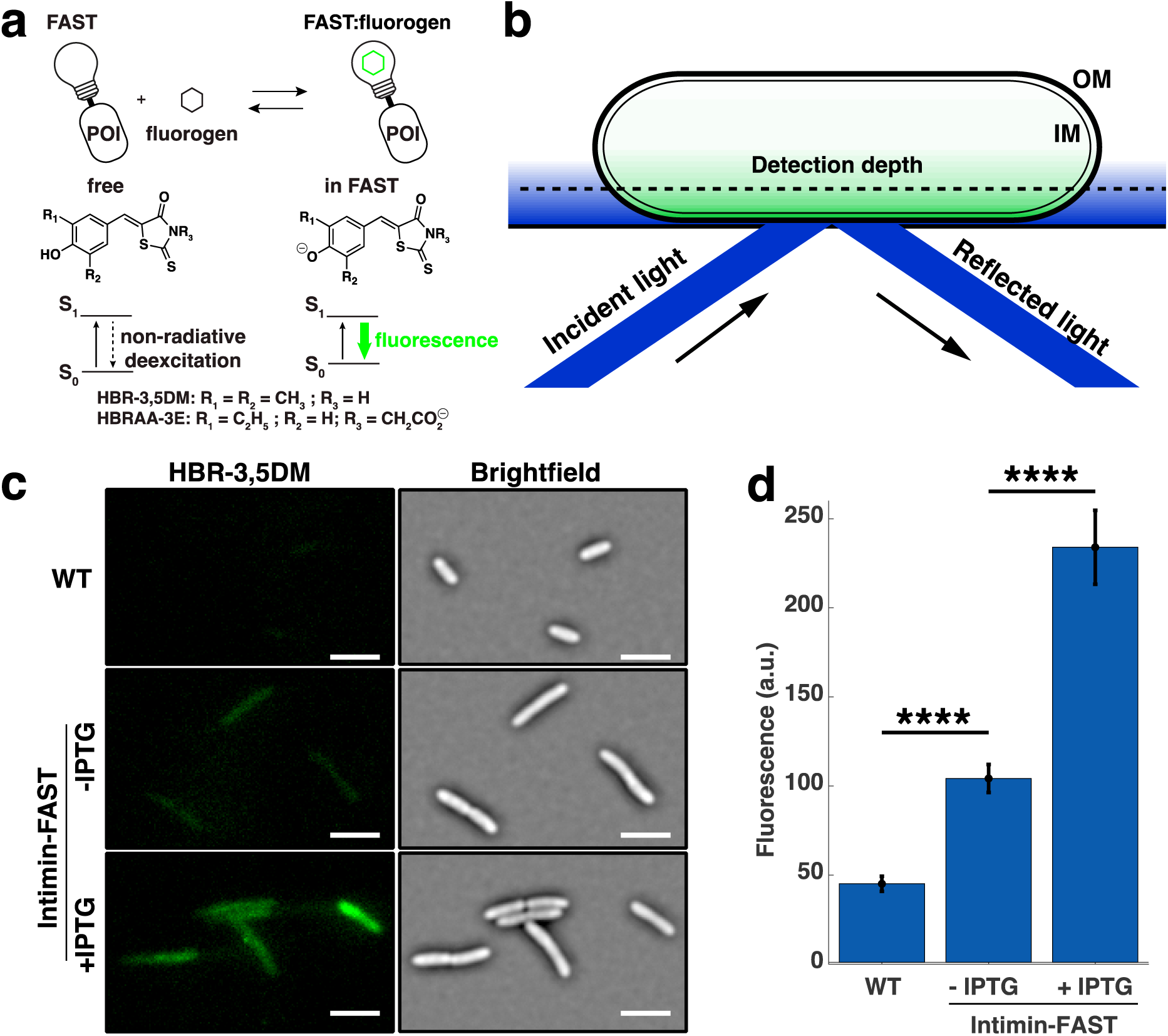
The chimeric Intimin-FAST protein expressed by *E. coli* cells becomes fluorescent upon binding to the membrane permeant fluorogen HBR-3,5DM. **a**. FAST is fused to a protein of interest (POI). The fluorogen HBR-3,5DM supplied in the culture medium becomes fluorescent when it binds to FAST and reveals the POI. **b**. Total internal reflection fluorescence microscopy (TIRF) generates an evanescent wave at the glass-gel interface in the sample. The short penetration depth of the evanescent wave (typically 200nm) allows to remove the unwanted background but only reveals a section of bacteria close to the coverslip. IM and OM are the inner and outer membrane respectively. **c**. Brightfield (right) and TIRF (left) images of *E. coli* MG1655 wild-type (WT) and Intimin-FAST cells. Exponentially growing cells (OD_590_=0.3) WT or containing pNeae2-FAST were deposited on a 1% agarose LB pad containing 20 µM of HBR-3,5DM with no (first two rows) or with 0.5 mM IPTG (bottom row). The background of the fluorescence images has been subtracted. All fluorescent images are displayed with the same contrast. Scale bar is 5 µm. **d**. Quantification of the fluorescent signal for WT and intimin-FAST cells in presence or absence of IPTG (n=152 for WT, n=108 for intimin-FAST with no IPTG, n=99 for intimin-FAST with 0.5 mM IPTG). Error bars represent standard errors.

Since the fluorogen HBR-3,5DM used to reveal FAST constructs permeated the cell membranes, TIRF microscopy did not guarantee that the fluorescent FAST-labeled intimin ß-domain was actually exposed on the outer membrane. Although FAST levels were 5 times higher than the non-specific background of the WT strains (Fig. 3d), we could not assert whether the functional FAST detected by the fluorescence of the fluorogen HBR-3,5DM actually corresponded to the cell-surface exposed intimin-FAST or to a fraction of the intimin-FAST located in the periplasm or in the cytoplasm.

### A non-permeant fluorogen to differentiate FAST localized in different *E. coli* cellular compartments

To address whether the signal observed with HBR-3,5DM was actually located at the cell-surface, we used a modified fluorogen, HBRAA-3E, that was previously reported not to permeate through the membranes of eukaryotic cells ^18^ but which capacity to cross bacterial membranes is unknown. We first checked if the non-permeant fluorogen HBRAA-3E did not stain the bacterial cytoplasm of *E. coli* using a strain constitutively expressing a free cytoplasmic FAST (Fig. 1b & 1e). While, by using TIRF microscopy, we detected FAST produced in the cytoplasm of *E. coli* as a uniform signal with the permeant fluorogen (HBR-3,5DM), no signal was detected when using the non-permeant fluorogen (HBRAA-3E) (Fig. 4a). This indicated that the non-permeant fluorogen was not able to cross the cytoplasmic inner membrane, but it does not necessarily mean that it might not cross the outer membrane of *E. coli*.

**Figure 4.**
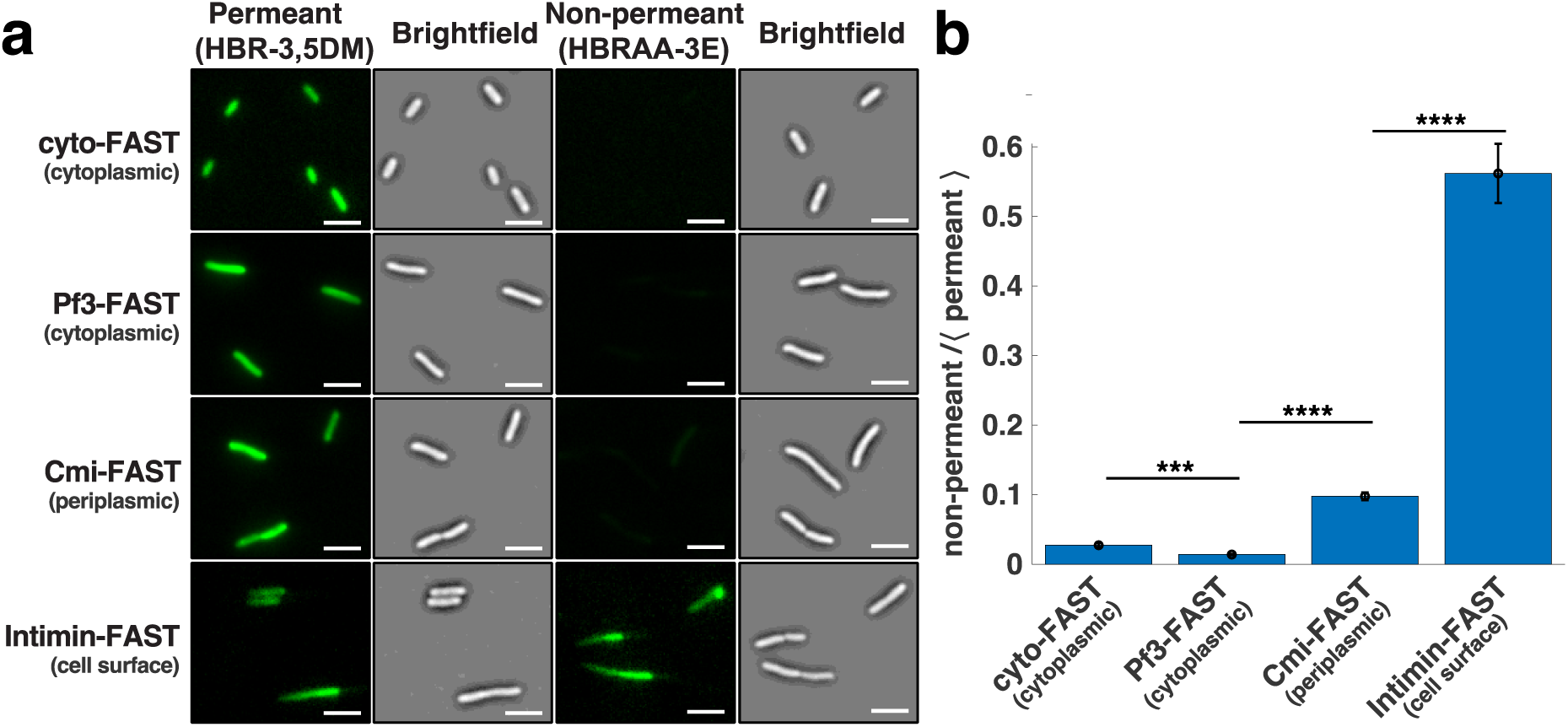
The outer membrane chimeric protein Intimin-FAST becomes fluorescent upon binding to the non-membrane permeant fluorogen HBRAA-3E. **a**. Brightfield (right) and TIRF (left) images of exponentially growing (OD_590_ = 0.3) *E. coli* harbouring FAST constructs revealed by the permeant (HBR-3,5DM) and non-permeant (HBRAA-3E) fluorogens. From top to bottom: cyto-FAST constitutively produced freely in the cytoplasm from the plasmid pZE1R-FAST; Pf3-FAST exposed on the cytoplasmic leaflet of the inner membrane produced from the plasmid pPf3-FAST; Cmi-FAST exposed on the periplasmic leaflet of the inner membrane produced from the plasmid pCmi-FAST; and intimin-FAST exposed at the cell surface produced from pNeae2-FAST, all in presence of IPTG. Cells were deposited on a 1% agarose LB pad containing either 20 µM of HBR-3,5DM or 40 µM of HBRAA-3E with 0.5 mM IPTG when necessary. The background of the fluorescence images has been subtracted. For each construct, fluorescent images obtained with the permeant and non-permeant fluorogens are displayed with the same contrast. Scale bar is 5 µm. **b** The signal of the non-permeant (HBRAA-3E) fluorogen divided by the average of the permeant (HBR-3,5DM) fluorescent signal for the different constructs: cyto-FAST (n=318 for HBR-3,5DM; n=55 for HBRAA-3E); Pf3-FAST (n=211 for HBR-3,5DM; N=198 for HBRAA-3E); Cmi-FAST (n=214 for HBR-3,5DM; n=160 for HBRAA-3) and intimin-FAST (n=99 for HBR-3,5DM; n=89 for HBRAA-3E). Error bars represent standard errors.

To further evaluate whether HBRAA-3E could cross the *E. coli* outer membrane, we built two strains expressing inner membrane-anchored protein either exposing FAST on the periplasmic, or on the cytoplasmic side of the inner membrane (Fig. 1c, 1d & 1e). In the first one FAST was fused at the C-terminus of the transmembrane domain of the major coat protein of the bacteriophage Pf3. The Pf3 transmembrane domain allows insertion into the inner membrane of bacteria with the protein facing the cytoplasm ^23–25^. In the second construction, FAST was fused at the C-terminus of the transmembrane domain of the specific immunity protein Cmi, which inhibits the action of colicin M in the periplasm of *E. coli* ^26^. The transmembrane domain of Cmi also allows insertion into the inner membrane, but unlike Pf3, Cmi exposes the residues fused to its C-terminus in the periplasm ^26^. Western blot on whole cell extracts using anti-E-tag primary antibodies demonstrated that both proteins were produced (Supplementary Fig. S2a). Both chimeric proteins migrated with an apparent higher molecular weight (MW) than expected (35 kDa instead of 24 kDa for Pf3-FAST; 30 kDa instead of 22 kDa for Cmi-FAST), possibly due to the hydrophobic nature of the transmembrane domain of the proteins. Consistent with antibodies inability to cross intact outer membranes, we detected no Cmi-FAST or Pf3-FAST immunofluorescence signal with anti-E-tag antibodies, in contrast with the surface-exposed Intimin-FAST (Supplementary Fig. S2b). We then added the membrane permeant fluorogen (HBR-3,5DM), and detected FAST signals for strains producing both Cmi-FAST and Pf3-FAST, indicating that these constructs were functional. On the contrary, we did not detect any periplasmic or cytoplasmic signals with the membrane non-permeant fluorogen (HBRAA-3E) in strains expressing Cmi-FAST or Pf3-FAST (Fig. 4a) indicating that HBRAA-3E was indeed a non-permeant fluorogen that could not cross the *E. coli* outer membrane in significant amount.

Finally, we used TIRF microscopy and both permeant and non-permeant fluorogens to probe the localization of functional intimin-FAST chimera and detected FAST signals with both fluorogens (Fig. 4a). The permeant fluorogen HBR-3,5DM revealed the total amount of FAST-tagged proteins expressed by cells, while the non-permeant fluorogen HBRAA-3E only revealed the fraction that was exposed on *E. coli* cell-surface. Since the two fluorogens have different emission/absorption spectra, brightness and affinity for the FAST amino-acid sequence (see methods), we could not quantitatively determine the absolute fraction of tagged proteins at the surface of cells. However, we could qualitatively compare strains by measuring the ratio between the non-permeant and the permeant signal. For the intimin-FAST construct the signal detected by the non-permeant fluorogen represented more than 50% of the total FAST revealed by the permeant fluorogen. For comparison this ratio was only 10% for the periplasmic Cmi-FAST construct and less than 2% for cytoplasmic constructs cyto-FAST and Pf3-FAST (Fig. 4b). These results confirmed that the non-permeant HBRAA-3E barely penetrates the outer membrane and that a large fraction of fluorescently labeled intimin-FAST is located on the cell-surface of *E. coli*. Besides, when bacteria are exponentially growing, a fraction of intimin-FAST fusions is located in the periplasmic space where the intimin protein undergoes folding by the SurA pathway ^27,28^. On the contrary, when bacteria are resuming growth from stationary phase, proteins must first be expressed, transported into the periplasm and folded before being exposed on the external face of the outer membrane. Consistently, when bacteria were harvested from stationary phase prior to observations, the appearance of the intimin-FAST signal revealed by a non-permeant fluorogen was delayed (Supplementary Fig. S3, Supplementary Movies S1 & S2). Interestingly, we noticed that the subcellular distribution of intimin-FAST fusions, which is homogeneous in exponential phase (Supplementary Movies S1 & S2), became polar when entering stationary phase (Supplementary Fig. S4, Supplementary Movies S3 & S4). This is consistent with previous studies using the SpyTag/SpyCatcher system ^13^.

### A fully functional FAST can also be exposed on the surface of gram-positive bacteria

Internalin B (InlB) is a surface protein involved in the invasion of host cells by the pathogenic monoderm bacterium *L. monocytogenes* ^29^. The association of InlB with the bacterial surface is labile ^30,31^. It has been reported to be mediated by affinity between the C-terminal GW domains of InlB with lipoteichoic acids in the bacterial membrane. Due to this non-covalent surface attachment, InlB has also been detected in soluble form in the culture medium of *L. monocytogenes* ^31^, which has been suggested to contribute to the activation of the InlB cell surface receptor, Met, at a distance from bacteria ^32^. Whereas the expression of *inlB* by *in-vitro* grown bacteria is low, it is activated *in vivo* by the main regulator of *L. monocytogenes* virulence genes, PrfA ^33^.

In order to evaluate whether FAST can be used to detect InlB, either when surface-exposed or when diffusible, the FAST coding sequence was introduced between InlB residues 342 and 343 in a flexible loop between the β2 and β3 leaflets of the B-repeat domain of the protein, that is neither involved in surface anchoring, nor in binding the Met receptor (Fig. 5a). This loop was selected, with the aim of minimizing the impact of introducing the FAST sequence on InlB structure and properties, because its sequence was less conserved among *L. monocytogenes* strains than the rest of InlB. The production of the InlB-FAST protein and its localization were assessed by immunodetection. We used an anti-InlB antiserum after fractionation of *L. monocytogenes* grown in Brain Heart Infusion (BHI) to separate proteins that were secreted and released in the medium (supernatant fraction SN), secreted proteins that were associated with the cell wall (cell wall fraction CW), proteins that were associated with the cytoplasmic membrane (membrane fraction MB, where InlB has been reported to be anchored) and cytoplasmic proteins (cytoplasm fraction CY) (Fig. 5b). Both InlB and InlB-FAST were almost exclusively detected in the MB fraction, indicating that InlB-FAST was exposed on the bacterial membrane in the same proportions as the wild-type protein, thus the introduction of FAST did not interfere with the anchoring of InlB to surface lipoteichoic acids.

**Figure 5.**
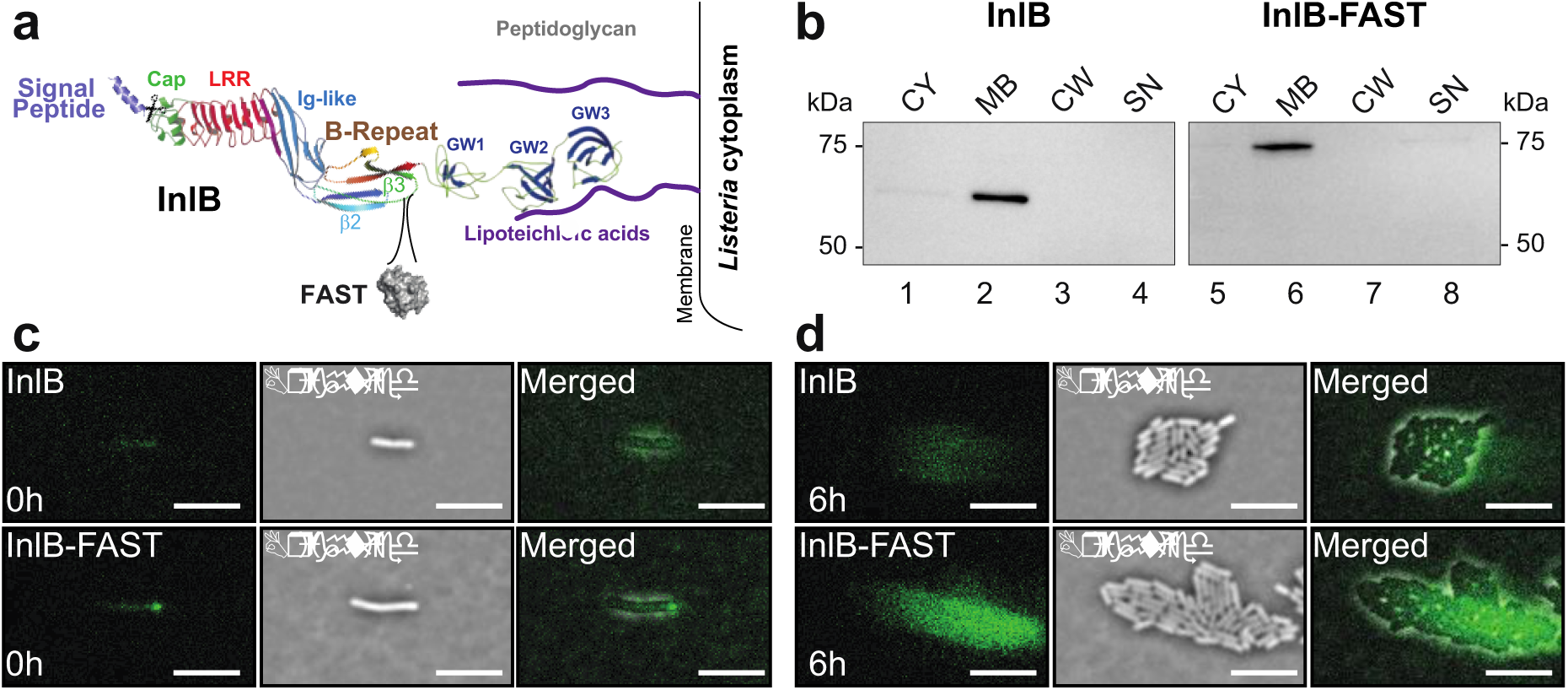
Relocation of InlB-FAST from stationary to exponential phase. **a.** Diagram of the InlB-FAST construct anchored to the membrane of *L. monocytogenes*. The insertion position of FAST, between the leaflets β2 and β3 of the “B-repeat” domain is indicated in orange. **b.** Western blot against InlB proteins for wild-type (WT) and InlB-FAST *L. monocytogenes*. CY = cytoplasmic fraction; MB = membrane fraction; CW = wall fraction; SN = supernatant fraction. In **c, d:** *L. monocytogenes prfA*\* that produced either InlB or InlB-FAST were grown in BHI with 20 µM HBR-3,5DM and imaged by TIRF microscopy. The contrast is the same on all images. Cells were resuming from stationnary phase in **c** and were in exponential phase in **d**. The microcolonies shown in **d** have developed from the cells shown in **c**. Scale bar = 5 µm.

In order to observe the localization of InlB at the single cell level, a culture of *L. monocytogenes* carrying the *prfA\** mutation and producing InlB-FAST was observed in TIRF microscopy. The *prfA*\* allele encodes a PrfA variant with a G145S substitution that is constitutively active and thereby leads to the high expression of PrfA-dependent virulence genes, including that of *inlB* ^34^. This strategy allowed us to bypass the need for PrfA activation that is otherwise triggered *in vivo*. Bacteria issued from a stationary phase were deposited on a nutrient BHI gel containing the permeant fluorogen HBR-3,5DM. When resuming growth, the location of InlB-FAST fusion was polar for 28% of cells (n=184) (Fig. 5c). However, the distribution of InlB became more diffuse as the microcolony grew, possibly due to labile binding that allows the release of InlB molecules from the bacterial surface as previously reported ^30,31^ (Fig. 5d, Supplementary Movie S5). On the contrary, the non-tagged strain showed a constant non-specific signal demonstrating that FAST remained fully functional when exported outside of *L. monocytogenes* cell-wall (Fig. 5c & d, Supplementary Fig. S5). Our results are consistent with previous immunofluorescence observations showing that the location of InlA, another internalin involved in *Listeria* invasion of host cells, redistributes from the lateral side to poles in stationary phase ^35^.

As a conclusion, we have shown that the FAST reporter system efficiently tags proteins exposed at the surface of both gram-negative and gram-positive bacteria. We used this reporter system to tag the β-domain of *E. coli* intimin and InlB from *L. monocytogenes*, two bacterial surface proteins involved in adhesion to or invasion of host cells. In agreement with previous studies, we observed that these two proteins were distributed on the whole surface of rod-shaped bacteria during exponential growth, while becoming polar in stationary phase.

The use of FAST presents several benefits over the use of other strategies to label cell-surface proteins. FAST labeling requires off-the-shelf synthetic fluorogens and is almost instantaneous allowing the study of highly dynamic processes in real-time. The reversible binding of the fluorogen reduces bleaching issues, which is advantageous for long term observations. In addition, since the fluorogen undergoes a red shift in emission upon binding to FAST, it allows to discriminate the bound from the unbound fractions, which opens ways for live imaging without any washing steps. Furthermore, FAST offers the key possibility to exclusively label cell-surface proteins, or both surface-exposed and intracellular proteins, through the use of membrane-impermeant or membrane-permeant fluorogens, allowing higher versatility when studying the dynamics of protein exposition to the cell-surface.

## Methods

### Bacterial strains and culture conditions

*E. coli* K12 strain used in this study is MG1655 (F-,λ-, *rph-1)* obtained *E. coli* genetic stock center CGSC#6300. Plasmid construction were performed directly in this strain. All strains were grown in Lysogeny Broth (LB) (Corning) medium supplemented with the appropriate antibiotic for plasmid selection, at 37°C with shaking at 180 rpm. The concentrations of the different antibiotics used for *E. coli* are as follow: Chloramphenicol 25 µg/mL (Cm), Ampicillin 100 µg/mL (Amp).

*Listeria monocytogenes* strains used in this work were derived from the clinical isolate LL195 ^36^. The *prfA*\* mutant that carried a single nucleotide mutation in the *prfA* allele, encoding a PrfA variant with a G145S substitution ^34^, was previously generated in this background ^37^. Plasmid constructions were performed using chemically-competent *E. coli* NEB5α (New England Biolabs Cat# C2987). All strains were grown at 37°C, 180 rpm in brain heart infusion (BHI, Difco) for *L. monocytogenes*.

### Strains and plasmids constructions

In order to favor the expression of transgenes, the DNA coding sequence for FAST, were codon-optimized for *E. coli* and *L. monocytogenes* using the online Optimizer application (http://genomes.urv.es/OPTIMIZER/) in guided random mode. The optimized sequences were obtained as synthetic gene fragments (Epoch Life Science and Eurofins genomics, respectively) and can be provided upon request. For the constructions of pZE1R-FAST and pNeae2-FAST, the two vectors, pZE1R-GFP and pNeae2, were linearized by PCR and the FAST encoding gene was amplified in the same time using Phusion Flash High-Fidelity PCR Master-Mix (Thermo Scientific, F548). The linearization of pZE1R-GFP has been designed to remove the GFP coding sequence. The primers used to linearize the vectors and to amplify FAST-coding sequence carry at their ends 20 bp of homology to facilitate recombination during the Gibson reaction. To construct pPf3-FAST and pCmi-FAST we used the same strategy. The pNeae2 plasmid has been linearized by PCR and the Neae coding sequence has been removed. In the same time, the sequences coding Pf3, Cmi and FAST have been amplified by PCR. The primers used to amplify Pf3 and Cmi transmembrane domain carried, in one end, homology region with FAST in addition to the homology region with the vector at the other end. Before performing the Gibson reaction, the linearized vectors were treated with the Fast Digest DpnI enzyme (FD1704) for 30 min at 37°C to remove any remains of circular plasmid which served as a matrix in the first round of PCR. The reaction was stopped by incubating the mixture for 20 min at 80°C. The Gibson reaction was then performed as described in ^38^ using 2 µL of linearized plasmid (100 ng/µL), 2 µL of insert, 10 µL of Gibson Master Mix 2X(100μL 5X ISO Buffer, 0.2 μL 10,000 U/mL T5 exonuclease (NEB #M0363S), 6.25 μL 2,000 U/mL Phusion HF polymerase (NEB #M0530S), 50 μL 40,000 U/mL Taq DNA ligase (NEB #M0208S), 87 μL dH_2_O for 24 reactions) and q.s for 20µL of sterile H_2_O. The mix was then incubated for 30 min at 50°C and dialyzed for 30 min. 50 µL of competent bacteria were then transformed, by electroporation, with 2 µL of Gibson’s reaction product. After 1 h at 37°C in LB, bacteria were spread on LB supplemented with appropriate antibiotic.

For allelic replacement at the *inlB* locus, the shuttle vector pMAD ^39^ was used. A sequence encompassing 1,000 base pairs upstream and downstream the site of insertion of FAST in the B-repeats domain of *inlB* was amplified, between which the FAST sequence was introduced, flanked by two twelve base pairs linkers. These fragments were obtained by PCR on LL195 genomic DNA or on the pAD-FAST-Myc plasmid pBIRD15 ^37^, assembled by PCR and inserted at the SalI/BglII sites of the pMAD plasmid. Allelic replacements of the *inlB* open reading frame by this construct in the genomes of *L. monocytogenes* strains LL195 and LL195 *prfA*\* were obtained as previously described ^39^.

Primers used in this study are listed in Table S2.

### Digestion by the proteinase K

*E. coli* cells were inoculated from plate and shaken at 180 rpm overnight at 37°C in LB. Equivalent of 1 OD_600_ was centrifuged for 10 min at 1,500 x *g* at 4°C. The pellets were then resuspended in protease buffer (50mM Tris, pH 8.8). Then 0, 5, 10 or 15 µg.mL^-1^ of Proteinase K (Canvax) was added and the samples incubated for 15 min at 40°C. Proteinase K activity was then stopped with 5 mM PMSF (phenylmethylsulfonyl fluoride). The bacteria were then centrifuged for 10 min at 1,500 x *g* at 4°C, resuspended in 1 mL of 50mM Tris pH 8.8 buffer and used for western blot, immunofluorescence.

### Immunodetection using Western Blot

After proteinase K digestion, bacteria were pelleted, resuspended in 100 µL of 1X Laemli buffer (Bio-Rad #1610747) and boiled at 100°C for 10 min. The proteins of an 0.1 OD_600_ equivalent quantity of cells were separated by SDS-polyacrylamide denaturing gel electrophoresis using precast TGX 4-15% gradient gels (Bio-Rad). The proteins were then transferred to a 0,2µm nitrocellulose membrane using Trans-Blot Turbo RTA Transfer Kit, nitro (Bio-Rad) and Trans-blot Turbo Transfer System (Bio-Rad). In order to verify that all wells have been uniformly loaded and to ensure that the proteins have been transferred, the membrane was stained with ponceau red (0.2% Ponceau S in 3% trichloroacetic acid). The membranes were washed with PBST (Phosphate Buffered Saline with Tween: PBS 0.05% Tween20) and saturated with a solution of PBST + 5% skimmed milk powder for 1 h at room temperature. The membrane was washed twice in PBST before being incubated with the primary rabbit anti-E-tag antibody diluted 1/5,000 in PBST (Abcam, ab3397) for 1h at room temperature (RT). The membrane was saturated again with a solution of PBST + 5% milk for 1h at RT and washed twice in PBST before being incubated with a secondary goat HRP-linked anti-rabbit antibody diluted 1/10,000 in PBST (Abcam #98431) for 1h at RT. The membrane was then washed 4 times 15 min in PBST. The revelation was performed using an ECL kit (Amersham, ECL western blotting detection reagents) and the iBright™ CL1500 system (Thermofisher).

### Immunofluorescence

The microscopy black slides (12 wells) were washed successively with water, 70% EtOH and 100% EtOH and then quickly fired. The wells were treated with 0.1% Poly-L-Lysine (Sigma P8920) for 2 min and washed 3 times with 1 mL PBS 1X (Ozyme) and allowed to dry. After proteinase K digestion, bacteria were pelleted, washed twice with 1 mL PBS 1X and then resuspended in 1 mL PBS 1X. 50 μL of sample was placed in each well, left 5 min at RT for cell attachment and fixed with 4% paraformaldehyde (PFA) (Sigma P6148) for 10 min at RT. The PFA was removed and the wells washed 3 times with 1 mL PBS 1X, then the samples were quenched with 50 mM NH4Cl (in PBS 1X) for 3 min at RT, before 3 washes with 1 mL PBS and saturation with a 0.5% BSA solution in PBS 1X (Sigma A7888) for 15 min at RT. Excess of BSA was removed and the wells are covered with the primary rabbit 1/500e anti-E-tag antibody (Abcam, ab3397) in 0.5% PBS 1X-BSA for 45 min at RT. The wells were washed 3 times with 1 mL PBS 1X and covered with a mixture containing secondary goat anti-rabbit-Alexa488 antibodies (Invitrogen Molecular Probe 2mg.mL^-1^, 1/300e) and DAPI (1/100e, Invitrogen Molecular Probe 10 mg.mL^-1^ D1306) for 45 min at RT. The slides are then washed 3 times with 1 mL of PBS and once with 1 mL of water and dried. Lastly, slides were mounted in Dako fluorescent mounting medium (Dako S3023) and observed by epifluorescence microscopy.

### *L. monocytogenes* fractionation and immunoblots

Bacterial proteins were separated into four fractions: supernatant, cell wall, membrane and cytoplasm as follows (adapted from^31^). One milliliter of overnight culture of *L. monocytogenes* was pelleted by centrifugation for 30 s at 6,000 × *g*. The supernatant (SN) was passed through a 0.22 µm filter, and proteins were precipitated with 16 % TCA, overnight at 4°C. The precipitated proteins were recovered by centrifugation at 16,000 × *g*, 15 min, 4°C, then washed twice with 1 mL acetone and finally resuspended in 80 µl 1X Laemmli buffer (4% SDS, 50 mM Tris pH 6.8, 10% glycerol, 50 mM DTT, 0,01% Bromophenol blue).

The bacterial pellet was washed once in 2 mL PBS, once in 2 mL TMS buffer (10 mM Tris HCl pH 6.0, 10 mM MgCl2, 0.5 M sucrose), resuspended in 100 µL of TMS supplemented with 60 µg/mL mutanolysin and 2 mM serine protease inhibitor AEBSF, and incubated for 1 hour at 37°C with gentle agitation to digest the peptidoglycan cell wall. The protoplasts were recovered by centrifugation at 15,000 × *g* for 5 minutes. The supernatant corresponding to the cell wall fraction (CW was precipitated with 16% TCA as described above. The protoplasts were lysed in 200 µL of lysis buffer (100 mM Tris pH 7.5, 100 mM NaCl, 10 mM MgCl2) and disrupted by multiple freeze-thaw cycles. Membrane (MB) and cytoplasm (CY) fractions were separated by centrifugation at 21,000 × *g* for 15 min at 4°C. The pelleted MB fraction was resuspended in 200 µL of RIPA buffer (25 mM Tris HCl pH 7.6, 150 mM NaCl, 1% NP-40, 1% sodium deoxycholate, 0.1% SDS). All the collected fractions were supplemented with Laemmli buffer, heat denatured for 5 min at 95°C and each fraction were analyzed by sodium dodecyl sulfate-polyacrylamide gel electrophoresis (SDS-PAGE) and Western blotting.

For Western blot analysis, 10 µL of each fraction were separated on 4-15% Mini-Protean TGX gels (Bio-Rad) by SDS-PAGE, and transferred on a 0.1 µm Amersham Protran nitrocellulose membrane (GE Healthcare Cat #10600000) using a Pierce G2 Fast Blotter. Proteins were probed with anti-InlB mouse monoclonal antibody B4.6^30^ at a 1/1,000 dilution in PBS supplemented with 0.05% Tween-20 and 5% skimmed milk powder, followed by secondary hybridization with anti-Mouse IgG-heavy and light chain Antibody (Bethyl Cat# A90-116P, RRID:AB_67183) at a 1/50,000 dilution in the same buffer. Signals were detected using Pierce ECL Plus Western Blotting Substrate and a Las4000 imager (GE Healthcare).

### Description of the Fluorogens

The preparation of HBR-3,5DM ^40^ and HBRAA-3E ^18^ was previously described. HBR-3,5DM and HBRAA-3E are commercially available from The Twinkle Factory (the-twinkle-factory.com) under the name ^TF^Amber, and ^TF^Amber-NP, respectively. FAST binds HBR-3,5DM and HBRAA-3E with a *K*Ds of 0.08 and 1.3 µM, respectively. FAST: HBR-3,5DM and FAST: HBRAA-3E complexes are characterized by absorption/emission peaks of 499 / 562 nm and 505 / 559 nm, and molecular brightness (*i.e.* the product of the fluorescence quantum yield and the absorptivity) of 23,500 M^−^1.cm^−^1 and 5,000 M^-1^.cm^-1^, respectively.

### TIRF Microscopy

*E. coli* and *L. monocytogenes* strains were inoculated from glycerol stocks and grown overnight at 37°C with shaking at 180 rpm in lysogeny broth (LB) and brain heart infusion (BHI), respectively. The next day, cultures were diluted (10^3^ times) in fresh medium and seeded on a gel pad (1% agarose with culture medium: LB or BHI) after reaching OD_590_ = 0.3 for *E. coli* single cell measurements or directly from stationary phase for microcolonies experiments. The gel pads were loaded with either 20 µM of the permeant (HBR-3,5DM) or 40 µM of the non-permeant (HBRAA-3E) fluorogens. The preparation was sealed on a glass coverslip with double-sided tape (Gene Frame, Fischer Scientific). A duct was previously cut through the center of the pad to allow for oxygen diffusion into the gel. Temperature was maintained at 37°C using a custom-made temperature controller. Bacteria were imaged on a custom microscope using a 100X/NA 1.45 objective lens (UPlan-Apo, Olympus) and an iXON EMCCD camera (Andor). Fluorescent images were taken using objective-based TIRF and laser excitation (Calypso 491nm, Cobolt). We used an emission filter of 590±20nm and a dichroic mirror of 540±10nm (Semrock). Image acquisition and microscope control were actuated with a LabView interface (National Instruments). Brightfield images were reconstructed from a z-stack of brightfield images ^22^. Brightfield images were used to segment bacteria and compute the mean fluorescence per pixel in the mask of each bacteria.

### Image analysis

For single cell experiments, we analyzed 10 different fields of view in each condition. Fluorescence quantification was performed by segmenting cells using either brightfield or DAPI fluorescent images for TIRF or immunofluorescence experiments, respectively. For each cell, we computed the mean intensity per pixel by averaging the fluorescence on cell mask. The fluorescence of the background was computed on the complementary of cell masks and then subtracted from the fluorescent images. Single cell measurements for the permeant fluorogen were given as the mean and the standard errors in the population for each strain. We computed the average ratio between the non-permeant and permeant signals by dividing single cell measurements obtained with the non-permeant fluorogen by the average fluorescence obtained with the permeant fluorogen on the same strain (*i.e.* average fluorescence in the population measured in single cells for 10 different fields of view using the HBR-3,5DM fluorogen). For experiments on microcolonies, we segmented microcolonies and yielded a single fluorescent trace for each field of view. Since InlB is loosely bound to the cell surface and can diffuse in the gel, we subtracted the fluorescent background measured on the first frame on every fluorescent images of the *L. monocytogenes* InlB-FAST time lapse. Image and data analysis were performed using custom MatLab routines. Statistical analysis was performed using unpaired t-tests (*** = p-value<0.001, **** = p-value<0.0001).

## Acknowledgements

We thank the Institut Pierre-Gilles de Gennes for technical support and Marion Lemarignier for assistance with InlB constructs. Work in the groups of ND and CB was supported by the Agence Nationale pour la Recherche (ANR) ADOBE-PRC-2019, by an Institut Pasteur grant, by the French government’s Investissement d’Avenir Program, Laboratoire d’Excellence “Integrative Biology of Emerging Infectious Diseases” (grant n°ANR-10-LABX-62-IBEID) and the *Fondation pour la Recherche Médicale* (grant no. DEQ20180339185). Work in the group of AL has received support under the program “Investissements d’Avenir” implemented by ANR (ANR-10-LABX-54 MemoLife and ANR-10-IDEX-0001-02 PSL University), Fondation pour la Recherche Médicale (FRM-AJE20131128944), Inserm ATIP-Avenir and Mairie de Paris (programme Émergences – Recherche médicale). Work in the group of AG was supported by the European Research Council (ERC-2016-CoG-724705 FLUOSWITCH). YC was supported by a MENESR (Ministère Français de l’Education Nationale, de l’Enseignement Supérieur et de la Recherche) fellowship. CPC received a doctoral fellowship from programme Interface pour le Vivant from Sorbonne University. DD was funded a Marie Sklodowska-Curie Action MSCA-IF-2018 (ERC).

## Data Availability

The datasets generated during and/or analysed during the current study are available from the corresponding author on reasonable request.

## Author Contributions

YC, CPC and DDA conducted the experiments. CB, ND and AL supervised the work. CB, ND, AL, CL, AG and JFA contributed to the methodology development. YC, CB, ND, CPC, DDA, AL and JMG analyzed the data. YC, CB, ND and AL wrote the manuscript with significant help from CPC, DDA, JMG and AG. All authors read and edited the manuscript.

