## Supplementary Figures and Tables for "Visualizing the dynamics of exported bacterial proteins with the chemogenetic fluorescent reporter FAST"

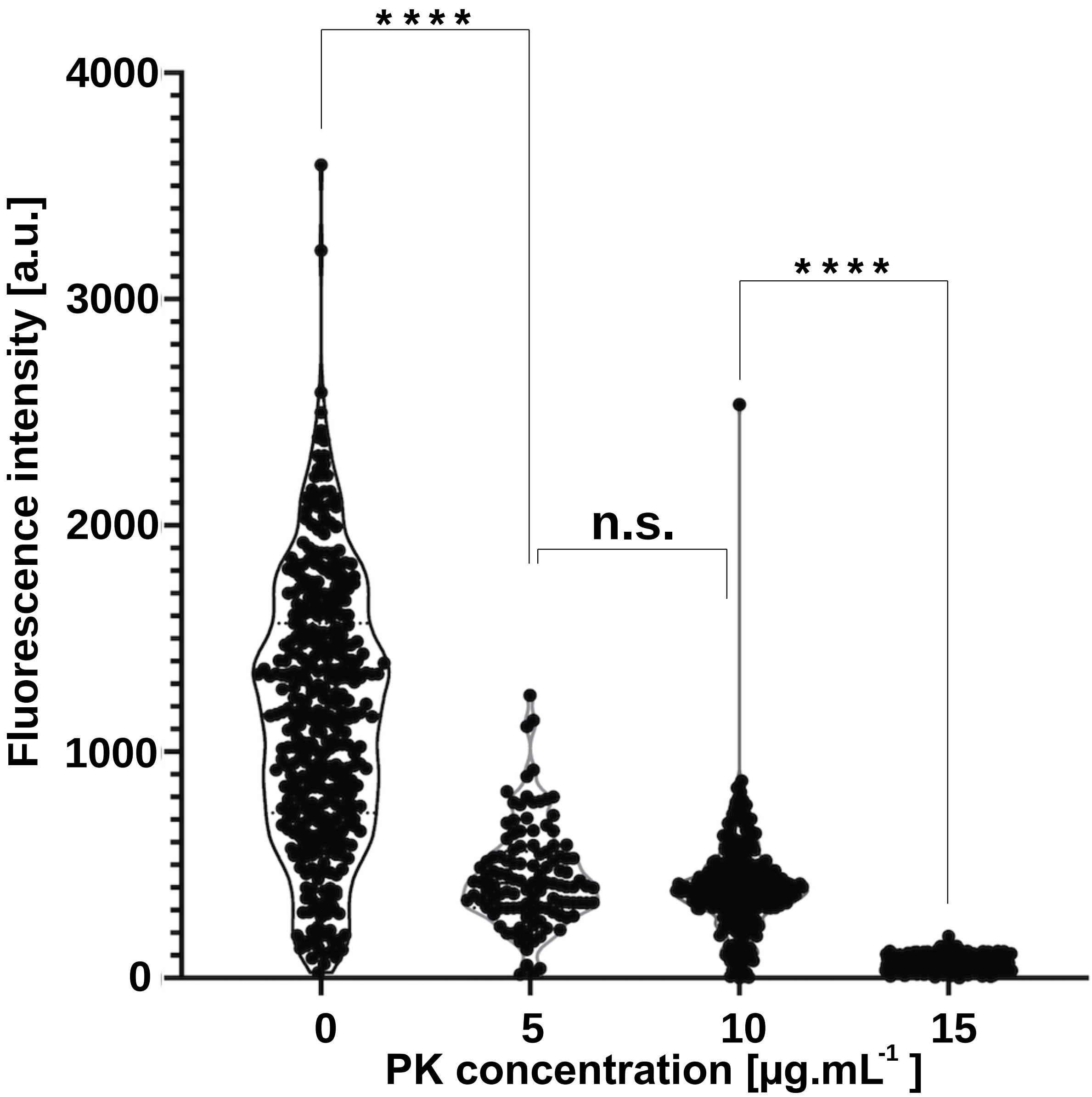


**Figure S1. Proteinase K degrades surface exposed FAST.** Quantification of fluorescence intensity from immunofluorescence images generated in Figure 2B. The distributions of fluorescence intensities are represented through violin plots for four values of proteinase K (PK) concentration. Data point sample size: n = (188, 143, 192, 78) for (0, 5, 10, 15) µg.mL$_{-1}$ proteinase K concentration.

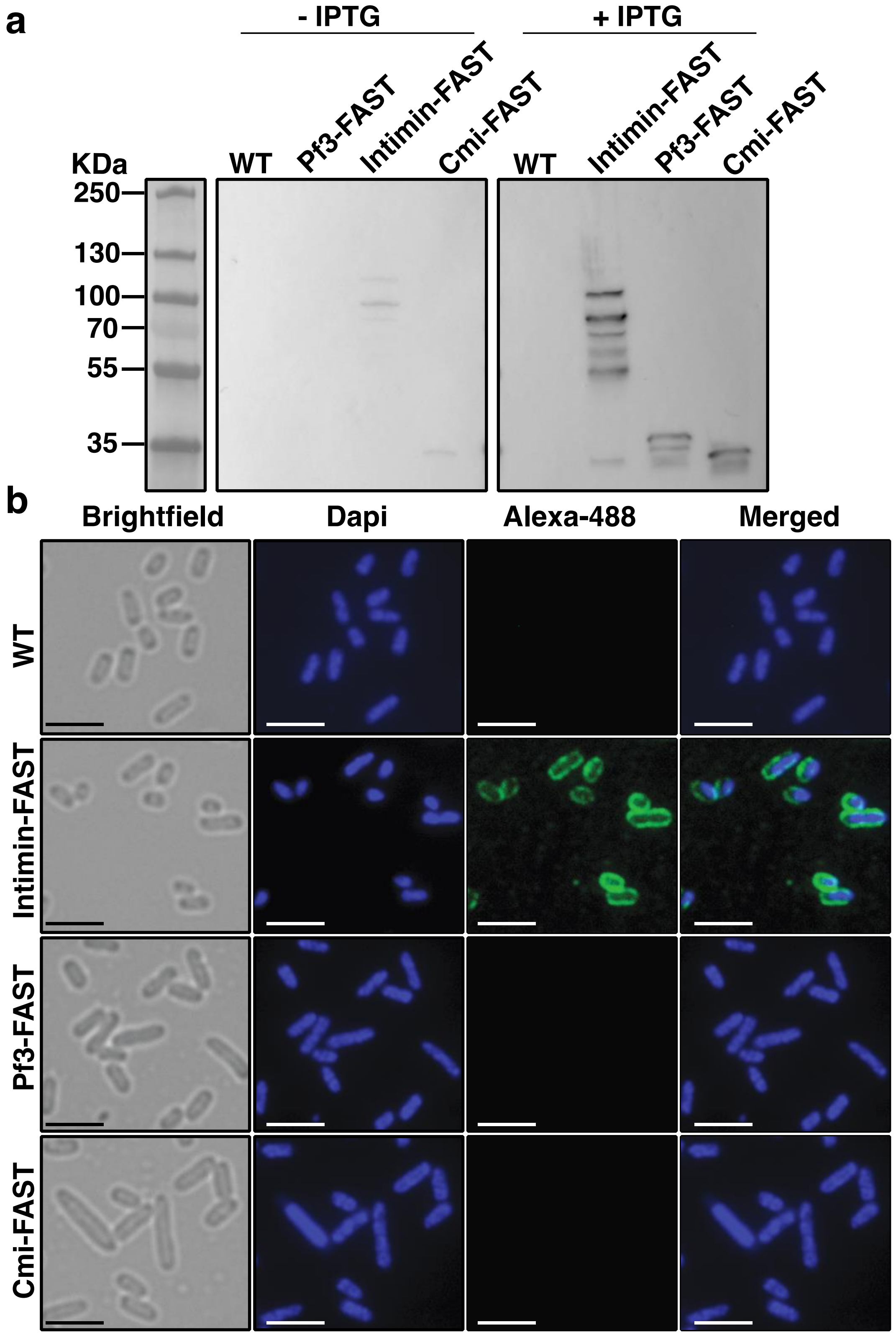


**Figure S2. Pf3 and Cmi-FAST fusions are produced by bacteria and are not detectable by immunofluorescence confirming their cytoplasmic/periplasmic localization. a :** Western blot of whole-cell protein extracts from *E. coli* MG1655, either uninduced (left pannel) or induced with 0.25 mM IPTG until OD 1 (right pannel), harbouring either no plasmid, pNeae2-FAST, pPf3-FAST or pCmi-FAST. **b:** Immunofluorescence performed on the corresponding cells, without permeabilisation. The bacterial surface was labeled with an anti-E-tag primary antibody and an anti-rabbit Alexa 488 conjugated secondary antibody. Scale bars = 2 µm.

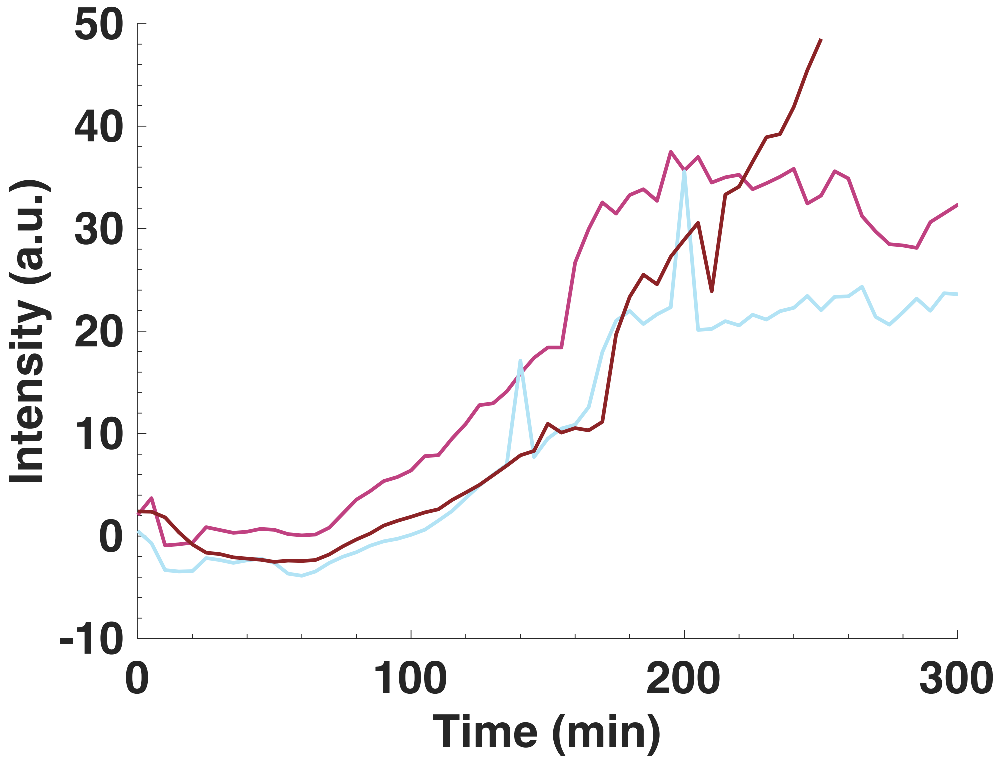


**Figure S3. Dynamics of intimin-FAST in microcolonies of *E. coli*.** Average intensity in microcolonies as a function of time for cells grown in LB supplemented with 40µM HBRAA-3E and IPTG at 0.5 mM for production of initimin-FAST from the plasmid pNea2-FAST. Each trace corresponds to an independent experiment.

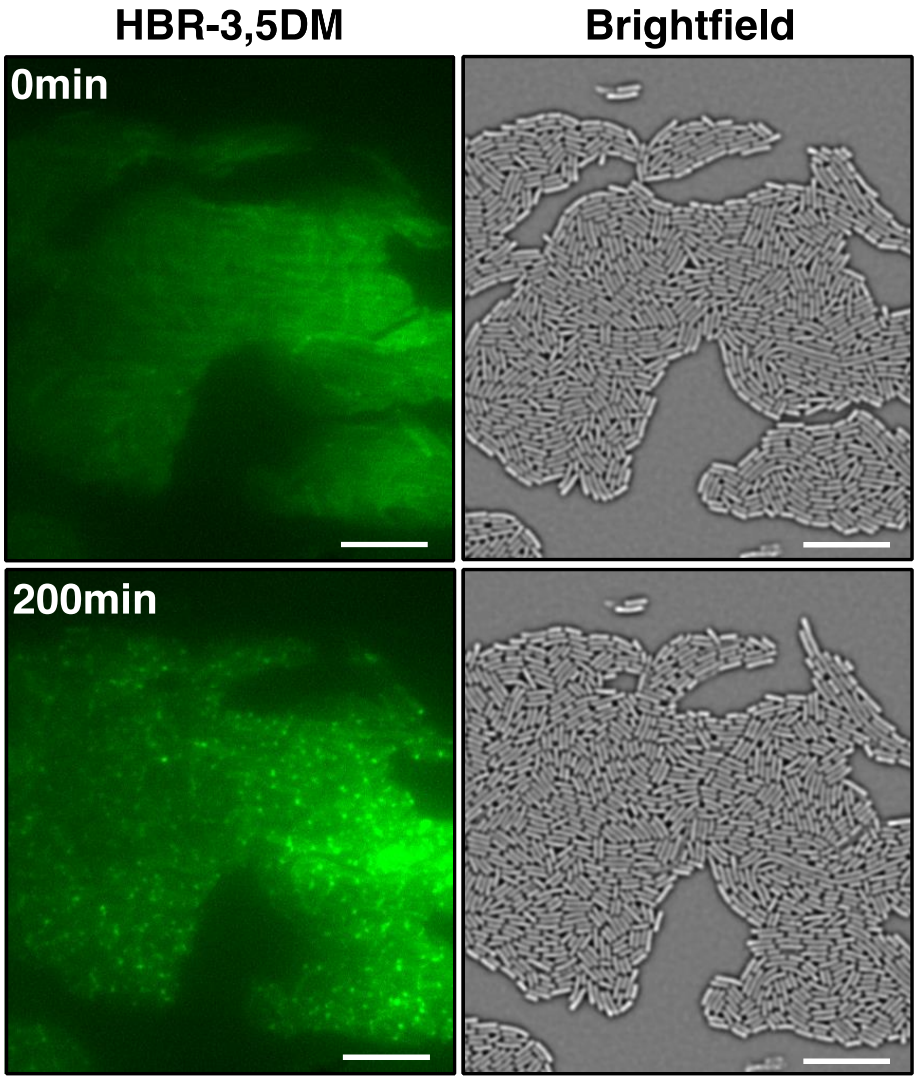


**Figure S4. The intimin-FAST chimeric protein is relocalized at cell pole in stationary phase.** TIRF and brightfield images of a microcolony of *E. coli* producing intimin-FAST, before (0min) and after (200min) the onset of stationary phase. The gel was loaded with 20 µM HBR-3,5DM and 0.5 mM IPTG. Scale bar = 10 µm.

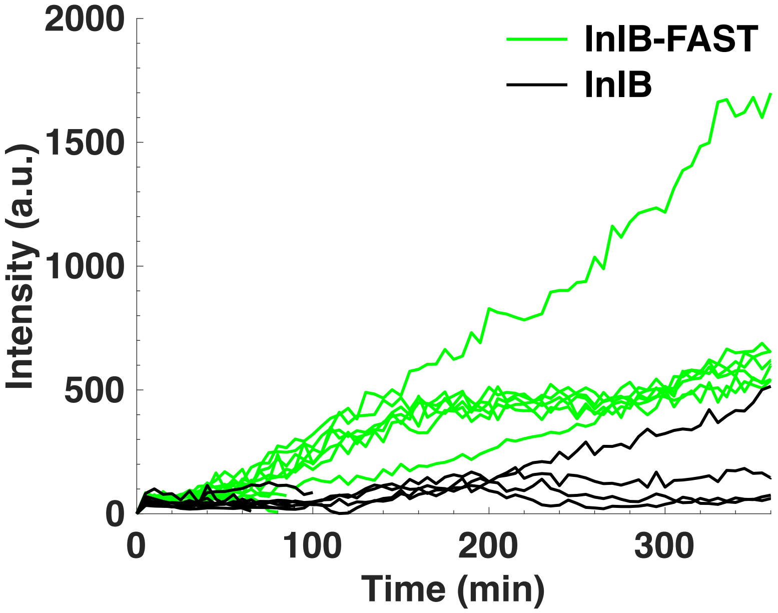


**Figure S5. Dynamics of InlB-FAST in microcolonies of *L. monocytogenes*. A.** Average intensity in microcolonies as a function of time for InlB-FAST (green) and InlB wild-type (black) cells growing in BHI supplemented with 20µM HBR-3,5DM. Each trace corresponds to an independent experiment.

### Supplementary Movies

**Movie S1**: Time lapse TIRF microscopy of *E. coli* intimin-FAST microcolonies growing in LB supplemented with 40 µM HBRAA-3E. Bacteria were inoculated from a culture in stationary phase. The time between frames is 5min.

**Movie S2**: Time lapse brightfield microscopy of *E. coli* intimin-FAST microcolonies growing in LB supplemented with 40 µM HBRAA-3E. Bacteria were inoculated from a culture in stationary phase. The time between frames is 5min.

**Movie S3**: Time lapse TIRF microscopy of *E. coli* intimin-FAST microcolonies growing in LB supplemented with 20 µM HBR-3,5DM. Bacteria were inoculated from a culture in stationary phase. The time between frames is 5min.

**Movie S4**: Time lapse brightfield microscopy of *E. coli* intimin-FAST microcolonies growing in LB supplemented with 20 µM HBR-3,5DM. Bacteria were inoculated from a culture in stationary phase. The time between frames is 5min.

**Movie S5**: Time lapse of *L. monocytogenes* InlB-FAST microcolonies growing in BHI supplemented with 20 µM HBR-3,5DM. Brightfield and TIRF microscopy are superimposed. The contrast, of brightfield images has been inverted for clarity. Bacteria were inoculated from a culture in stationary phase. The time between frames is 5min.

### Supplementary Tables

| **Strains** | **Description** | **Source** |
|---|---|---|
| ***Escherichia coli*** | | |
| MG1655 | $F^-$,$\lambda^-$, *rph-1* | *E. coli* genetic stock center CGSC#6300. |
| ***Listeria monocytogenes*** | | |
| BIRD111 | *E. coli* NEB5α, pMAD-*$inlB_{1-1026}$-FAST-$inlB_{1027-end}$*, ($Amp_R$) | This study |
| BIRD119 | *L. monocytogenes* LL195, *inlB-FAST* | |
| BIRD 232 | *L. monocytogenes* LL195, *prfA*, inlB-FAST* | |
| BIRD 234 | *L. monocytogenes* LL195, *prfA** | 37 |
| **Plasmids** | | |
| pNeae2 | Plasmid containing N-terminal fragment of intimin with N-terminal signal peptide, periplasmic LysM domain, a β-barrel domain and the Ig-like domains (D00-D0). In the plasmid pNeae2 three different tags (*E-tag, His-tag* and myc-tag) have been fused in frame with the C-terminal end of the D0 domain. All is under control of the *pLac* promoter. CmR | 21 |
| pZE1R-GFP | Plasmid containing GFP under control of the phage λ *PcL* promoter. GFP is constitutively express in the cytoplasm of bacteria. AmpR | 41 |
| pPf3-FAST | Optimized FAST encoding gene cloned in pNeae2. Neae2 domain has been removed by PCR. FAST is fused to the Pf3 transmembrane domain. The construction is under control of the *plac* promoter. CmR | |
| pCmi-FAST | Optimized FAST encoding gene cloned in pNeae2. Neae2 | |

| | | |
|---|---|---|
| | domain has been removed by PCR. FAST is fused to the Cmi transmembrane domain. The construction is under control of the *plac* promoter. CmR | This study |
| pNeae2-FAST | Optimized FAST encoding gene cloned in pNeae2. FAST is fused to the $\beta$ domain of the intimin. The construction is under control of the *plac* promoter. CmR | |
| pZE1R-FAST | Optimized FAST encoding gene cloned in pZE1R-GFP. GFP has been removed by PCR. FAST is under control of the phage $\lambda$ *PcL* promoter and is constitutively express in the cytoplasm of bacteria. AmpR | |
| pMAD-*inlB*$_{1\text{-}1026}$- FAST-*inlB*$_{1027\text{-}end}$ (pBIRD111) | Plasmid for allelic replacement at the *InlB* locus: optimized *FAST* encoding gene has been inserted by PCR between the $\beta$2 and $\beta$3 leaflets of the B-repeat domain of the InlB protein (at position 1026 ± 1000 bp). ErmR (*Lm*) and AmpR (*Ec*) | |
| pAD-*FAST*-*Myc* (pBIRD15) | Integrative plasmid containing the optimized *FAST* encoding gene fused to a Myc-tag, under the P$_{HYPER}$ constitutive promoter. pAD is integrated at the tRNA$_{Arg}$ locus in the genome of *L. monocytogenes*. CmR | 37 |

**Table S1. Strains and plasmids used in this study**

| Name | Sequence |
|---|---|
| ***E. coli*** | |
| pZ-FAST.REV | cgtgttccatgcggtacctttctcctctttaatga |
| pZ-FAST.FOR | cgtgtttaataagcttaattagctgagtctagaggcat |
| F2_pZ_FAST.REV | agctaattaagcttattaaacacgtttaacgaaaacccagtaagagt |
| F2_pZ_FAST.FOR | aaggtaccgcatggaacacgttgcgttcg |
| F1_Neae2-FAST.FOR | aacgtgtttaataaaagcttgacctgtgaagtgaaaaat |
| F1_Neae2-FAST.REV | gctaccgctaccgctgccgctacctgcagctgcatcctcttctgaga |
| F2_Neae2-FAST.FOR | ggcagcggtagcggtagcggcagcatggaacacgttgcgttcg |
| F2_Neae2-FAST.REV | tcaagcttttattaaacacgtttaacgaaaacccagtaagagt |
| vec_Neae2_FAST_CMI.F | aaggcgaatttaacaacaacggtgcgccggtgc |
| vec_Neae2_FAST_CMI.R | ttcatgctaatcactttcatgatttgccctcgttatctagaaat |
| CMIforNeae2_FAST_CMI.F | ctagataacgagggcaaatcatgaaagtgattagcatgaaatttatttttattctgacca |
| CMIforNeae2_FAST_CMI.R | ggatacggcaccggcgcaccgttgttgttaaattcgcctttatcttcgc |
| Vec_Neae2_FAST_pf3.F | tctttgaatttaacaacaacggtgcgccggtgc |
| Vec_Neae2_FAST_pf3.R | tcagtaatcacggattgcatgatttgccctcgttatctagaaat |
| pf3for_Neae2_FAST_pf3.R | ggatacggcaccggcgcaccgttgttgttaaattcaaagaattgcgctttg |
| pf3for_Neae2_FAST_pf3.F | Ctagataacgagggcaaatcatgcaatccgtgattactgatgtgac |
| ***L . monocytogenes*** | |
| oAL833<br>(3' *inlB* LL195 (-1000 from position 1026) – BglII-pMAD, Fw) | gacagatctgaaggaaatattgtttaatctcagg |
| oAL834<br>(*inlB* LL195(to position 1026)-GGSG linker -3'*FAST*, Rv) | catgccgctcgagcccgtccctgcctctacttttg |
| oAL835<br>(*inlB* LL195(to position 1026)-GGSG linker-3'*FAST*, Fw) | gacgggctcgagcggcatggaacatgttgctttcggttc |
| oAL836<br>(5' *FAST*-AAAG linker-*inlB* from pos 1027, Rv) | tcggcccgcggccgctacacgtttaacaaaaaccc |
| oAL837<br>(5' *FAST*-AAAG linker-*inlB* from pos 1027, Fw) | gtagcggccgcgggccgaataactgcacctaaacc |
| oAL838<br>(5' *inlB* LL195(+1000 from pos 1026)-SalI-pMAD, Rv) | gacgtcgacccggccgcagaatatgg |

**Table S2. Primers used in this study**
